# *Ptpn14* Knockout Mice Reveal Critical Female-Specific Roles for the Hippo Pathway

**DOI:** 10.1101/2025.01.13.632464

**Authors:** Edel M. McCrea, Neoklis Makrides, Takako Tabata, Hannes Vogel, Brooke Howitt, Wencke Reineking, José G. Vilches-Moure, Mengxiong Wang, Xin Zhang, Laura D. Attardi

## Abstract

The Hippo pathway regulates many physiological processes, including development, tumor suppression, and wound healing. One understudied Hippo pathway component is PTPN14, an evolutionarily conserved tyrosine-phosphatase that inhibits YAP/TAZ. While an established tumor suppressor, PTPN14’s role in tissue homeostasis has remained unclear. We thus generated *Ptpn14*-deficient mice and found that only ∼60% of *Ptpn14^-/-^* mice survived postnatally, highlighting the importance of PTPN14 for viability, while also enabling the discovery of PTPN14 physiological functions. *Ptpn14^-/-^*mice developed debilitating corneal lesions and the uterus defect, hydrometra, as well as heart and kidney abnormalities. *Ptpn14*-deficiency precipitated an impaired injury response in the cornea and dysregulated YAP signaling in the uterus. Notably, these phenotypes were female-specific, revealing sexually-dimorphic Hippo pathway function through PTPN14. Finally, analysis of human *PTPN14* variants suggested that PTPN14’s essential roles are conserved in humans, underscoring the importance of our insights for designing therapies to improve women’s health.

## INTRODUCTION

The Hippo pathway, named originally for the overgrowth phenotype in Drosophila mutants, is a highly evolutionarily conserved signaling pathway that regulates cellular proliferation, survival, polarity, cell fate, and, at the macroscopic level, organ size^1^. Central to this pathway is a core kinase cascade, comprising the SAV1, MST1/2, MOB1A/B and LATS1/2 proteins, which responds to extracellular signals that ultimately converge on the YAP/TAZ transcriptional coactivator(s) to modulate specific gene expression programs. When the Hippo pathway is off, YAP/TAZ enter the nucleus and bind the sequence-specific transcription factor, TEAD, leading to the expression of genes promoting growth and survival. The Hippo pathway becomes activated in response to a variety of cues, including matrix stiffness, cell-cell contact, GPCR signaling, glucose metabolism, and hormone signaling, which trigger phosphorylation of SAV1, causing it to phosphorylate MST1/2, which then phosphorylates LATS1/2^2^. MOB1A/B acts as a protein scaffold for LATS1/2, which promotes its phosphorylation by MST1/2^3,4^. Once phosphorylated, LATS1/2 phosphorylates YAP/TAZ, which sequesters it in the cytoplasm and prevents it from activating growth-promoting gene expression programs^5–7^. The careful modulation of YAP/TAZ signaling by the Hippo pathway is essential for organismal health, as hyperactivation of YAP can result in cancer, oversized organs, and other deleterious effects, while YAP ablation causes severe developmental defects or lethality.

In addition to the core cascade, various other Hippo pathway proteins can fine-tune modulation of YAP/TAZ. These include the tumor suppressor, NF2, which promotes activation of LATS1/2 directly. AMOT, another protein outside the core kinase cascade, also can promote phosphorylation of LATS1/2 and/or sequester YAP in the cytoplasm at F-actin^8^. These examples represent a small number of the known Hippo pathway proteins, underscoring the great complexity of this signaling network. In fact, one study proposed a two-Hippo pathway model in which YAP/TAZ inhibition through MST1/2 and SAV1 regulated organ size to a greater extent than control of YAP by NF2 and MAP4Ks, which played a larger role in regulating the growth of monolayer cell linings and tubular structures^9^. Understanding the common and distinct contributions of individual Hippo pathway constituents is therefore of paramount importance for understanding their roles in physiology and pathology.

The use of conditional and constitutive knockout (KO) mouse models has illuminated various facets of Hippo signaling. Although each element of the Hippo pathway definitively contributes to YAP inhibition, the effect of knocking out each of these individual Hippo pathway components varies widely both within and across tissues. In numerous knockout mice deficient for Hippo pathway components, including *Mst1/2* and *Nf2*, prominent phenotypes arise in the liver, such as a dramatic increase in liver size and the development of hepatocellular carcinoma^10–14^. In contrast, knocking out other components of the Hippo pathway, such as AMOT, has no effect on liver development or function^15^. Further, knockout of *Mst1/2* in the mouse epidermis has no obvious phenotype, while deletion of *Mob1a/b* in the epidermis leads to tumorigenesis^16,17^. These data suggest that the many components of this network must be studied in multiple tissue types for us to fully understand this complex pathway.

Hippo pathway knockout mice have also revealed the importance of the pathway in injury repair in many tissues, including the heart, liver, intestine, skin, and nervous system^18^. For example, an intestine-specific YAP cKO caused no intestinal defects at homeostasis but prevented the intestines from healing following injury^19^. Analysis of mice lacking *Sav1* in cardiomyocytes showed greater heart regeneration than in wild-type counterparts in response to myocardial infarction, supporting a role for Hippo signaling in the context of regeneration. This result was replicated both in mice and pigs using an shRNA targeting SAV1, which was delivered to the injured heart via adenovirus^20,21^. This SAV1 gene therapy approach is under investigation for use in humans and underscores the utility of mouse models in understanding the Hippo pathway. Fully characterizing the role of the Hippo pathway in injury responses will enable the development of therapies facilitating tissue regeneration and enhanced wound healing in the future.

*Ptpn14*, another upstream member of the Hippo pathway, encodes a tyrosine phosphatase that was first discovered in a screen for tyrosine phosphatases expressed in normal human breast tissue and is highly evolutionarily conserved from *Drosophila* to humans^22^. Although a clear role for its phosphatase domain has yet to be established, PTPN14 uses its two internal PPxY binding sites to bring LATS1/2 and YAP/TAZ into closer proximity, thereby functioning as a protein scaffold to promote LATS1/2-dependent phosphorylation of YAP/TAZ^23^. Like other members of the Hippo pathway, which are generally classified as tumor suppressors due to their ability to restrain YAP/TAZ, PTPN14 is also a tumor suppressor, displaying high mutation rates in basal cell carcinoma, cervical cancer, lung cancer, cholangiocarcinoma, colorectal cancer, and other cancers^24–27^. While the role of PTPN14 as a tumor suppressor is widely documented, its roles in mammalian development and tissue homeostasis remain largely unknown. Inactivating mutations in the *Drosophila* homologue, *Pez*, resulted in delayed development, reduced body size, compromised fertility, and an accumulation of progenitor cells in the adult midgut, while overexpression of *Pez* in the midgut inhibited the increased proliferation of progenitor cells in response to injury, but had no effect on these cells at homeostasis^28^. In zebrafish, knockdown of *ptpn14* in embryos caused an overproliferation of the ventricular zone of the brain, malformed pharyngeal arches in the heart, and severe pericardial edema^29^. These data, taken together with the high sequence conservation of *PTPN14* across evolution, suggest that PTPN14 may contribute to important signaling pathways relating to development and normal tissue function in mammals. Thus, to elucidate the physiological roles of PTPN14 *in vivo*, we generated *Ptpn14* knockout mice and studied the phenotypes associated with *Ptpn14* deficiency at the organismal level in both males and females. Our investigation revealed multiple critical functions of PTPN14 in diverse organs, and, unexpectedly, a dramatic sex-bias for many of these phenotypes, thus illuminating the importance of this protein *in vivo*.

## RESULTS

### Generating Ptpn14 Conditional Knockout Mice

To understand the role of PTPN14 in development and tissue homeostasis, we generated *Ptpn14* conditional knockout mice using Cre-LoxP technology. In the design of the conditional knockout allele, we chose to flank exon 3 with LoxP sites, as deletion of exon 3 would shift the remainder of the protein out of frame (Figure 1A). After obtaining founders (see Materials and Methods), we performed multiple diagnostic PCRs to identify mice carrying the *Ptpn14* conditional knockout allele before expanding the colony. First, we confirmed the presence of the 5’ and 3’ LoxP sites on either side of exon 3 by PCR (Supplemental Figure 1A). We then verified integration into the correct location in the genome by designing 5’ and 3’ external PCRs in which one primer was located outside the region corresponding to the donor DNA used during the CRISPR targeting of the locus (Supplemental Figure 1A). In addition to the conditional knockout mice, we also generated constitutive *Ptpn14* knockout mice by crossing *Ptpn14^fl/fl^* mice with C57BL/6J *Deleter-Cre* mice. The *Deleter-Cre* transgene drives recombination in the germline, resulting in *Ptpn14* null alleles. After breeding out the Cre transgene, we propagated the colony and confirmed genetic recombination at the *Ptpn14* locus in both *Ptpn14^+/-^* and *Ptpn14^-/-^* mice (Supplemental Figure 1B).

**Figure 1.**
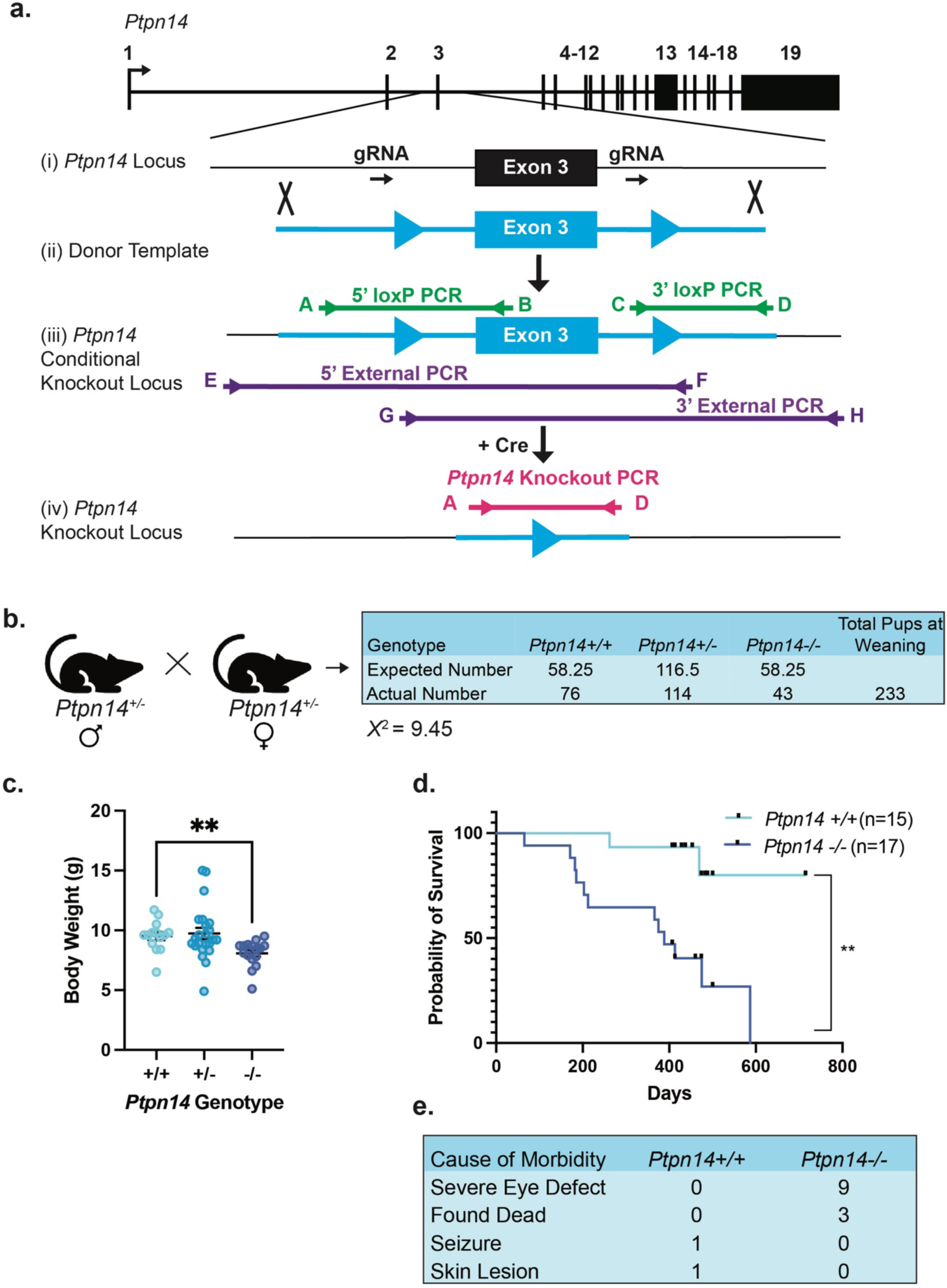
*Ptpn14 Deficiency* Compromises Viability. (A) Strategy for generating *Ptpn14* conditional and complete knockout strains. (i) *Ptpn14* locus with magnified view of exon 3. sgRNAs flanking exon 3 and (ii) donor DNA template containing floxed exon 3 are indicated. (iii) Initial targeting results in a *Ptpn14* conditional knockout allele. (iv) Upon expression of the Cre recombinase, Cre recombines the LoxP sites surrounding exon 3, leading to exon 3 deletion from the genome and a complete *Ptpn14* knockout allele. Sites for PCR primers (A-H) used to confirm the allele (see text) are highlighted. (B) Expected and actual pup numbers obtained at weaning after intercrossing *Ptpn14^+/-^* mice. Chi-square analysis indicates a significant disenrichment of *Ptpn14^+/-^* and *Ptpn14*^-/-^ mice (X^2^ = 9.45, the threshold for p-value < 0.05 is 5.99 for two degrees of freedom). (C) Body weight of *Ptpn14^+/+^* (n = 14), *Ptpn14^+/-^* (n = 23), and *Ptpn14^-/-^* (n = 16) mice at weaning (Day 20-25). (D) Kaplan- Meier curve showing the survival of *Ptpn14^+/+^* (n= 15) and *Ptpn14^-/-^* (n = 17) mice. (E) Phenotypes observed in the cohorts used in the Kaplan-Meier aging study in (D). One wild-type mouse died from a seizure, and one developed a skin infection that necessitated sacrifice.

To verify that Cre-mediated recombination of exon 3 resulted in loss of the PTPN14 protein, we intercrossed *Ptpn14^+/fl^* mice and made mouse embryonic fibroblasts (MEFs) at E13.5. After delivering adenovirus-Cre (Ad-Cre) or control empty adenovirus (Ad-Empty) to *Ptpn14^+/+^, Ptpn14^+/fl^,* and *Ptpn14^fl/fl^* MEFs, we found that only Ad-Cre transduction into the *Ptpn14^fl/fl^* MEFs caused loss of the PTPN14 protein (Supplemental Figure 1C). These findings confirmed that recombination of the *Ptpn14* conditional allele generates a *Ptpn14* null allele after Cre introduction.

### Ptpn14^-/-^ Mice Exhibit Premature Mortality and Complications with Breeding

To evaluate whether PTPN14 deficiency affects organismal viability, we intercrossed *Ptpn14^+/-^* mice and quantified the genotypes of their progeny at weaning (Figure 1B). Although a number of *Ptpn14^-/-^* mice survived to adulthood, Chi-squared analysis indicated there were significantly fewer *Ptpn14^-/-^* and *Ptpn14^+/-^* than expected by Mendelian ratios (Figure 1B). *Ptpn14^-/-^* mice also displayed a lower body weight at weaning, a sign that *Ptpn14* deficiency may compromise developmental processes and/or homeostasis (Figure 1C). Upon tracking the success of the breeding pairs in our colony, we found that out of 16 *Ptpn14^+/-^*female breeders, 2 females reached morbidity due to dystocia, or an inability push the pups out of the birthing canal. We also bred two *Ptpn14^-/-^* female mice, which did not successfully reproduce, with one carrying a single litter before succumbing to dystocia and the other never becoming pregnant at all. These observations of *Ptpn14^+/-^*and *Ptpn14^-/-^* mice suggest a negative effect on reproduction with PTPN14 loss.

Because many *Ptpn14^-/-^* mice reached adulthood, we aged these mice and assessed whether *Ptpn14* loss altered overall survival or caused any deleterious phenotypes. We generated cohorts of 15 wild-type mice (8 males and 7 females) and 17 *Ptpn14^-/-^* mice (7 males and 10 females) and measured their lifespan and any associated phenotypes upon morbidity. The *Ptpn14^-/-^*mice displayed reduced survival relative to wild-type counterparts (Figure 1D). Of the 17 *Ptpn14^-/-^* mice, three died prematurely and nine had severe eye defects necessitating sacrifice (Figure 1E). These defects manifested as large white protrusions emerging from the center of the eye. While no other external signs of poor health appeared in these knockout mice at morbidity, we observed other deleterious phenotypes, such as hydrometra and kidney cysts, after sacrifice (see below). These findings underscore the role of PTPN14 in normal tissue function.

### Ptpn14 Deficiency Dramatically Compromises Uterine Function

The dystocia observed in the *Ptpn14^-/-^* female mice suggested a potential defect in the female reproductive system. Indeed, at morbidity, all *Ptpn14^-/-^* female mice from the aging cohort and four mice sacrificed from the colony, all of which were virgins, exhibited distended uterine horns filled with fluid upon dissection, a condition known as hydrometra (n = 12, Figure 2A). The material filling the uterine horns was usually red, although in some mice it was white and cloudy, and upon histological evaluation the liquid appeared to be a mixture of proteinaceous material and blood (Figure 2B, C). While the wall of the uterine horns was severely distended to accommodate the liquid accumulating there, no other abnormalities were detected in the uterine wall musculature (Figure 2D), suggesting that the liquid might be trapped there by an obstruction further down in the uterus or the cervix. We also noted severely dilated blood vessels located within the uterine wall that could have resulted from blocked blood flow, further suggesting an obstruction occurring somewhere in the organ (Figure 2E). While hydrometra is typically caused by a blockage of the female reproductive tract, histological evaluation failed to detect the presence of an obstruction in the uterus, cervix, or vagina. However, these membranes can rupture immediately upon dissection, and therefore the cause of hydrometra can often not be definitively determined.

**Figure 2.**
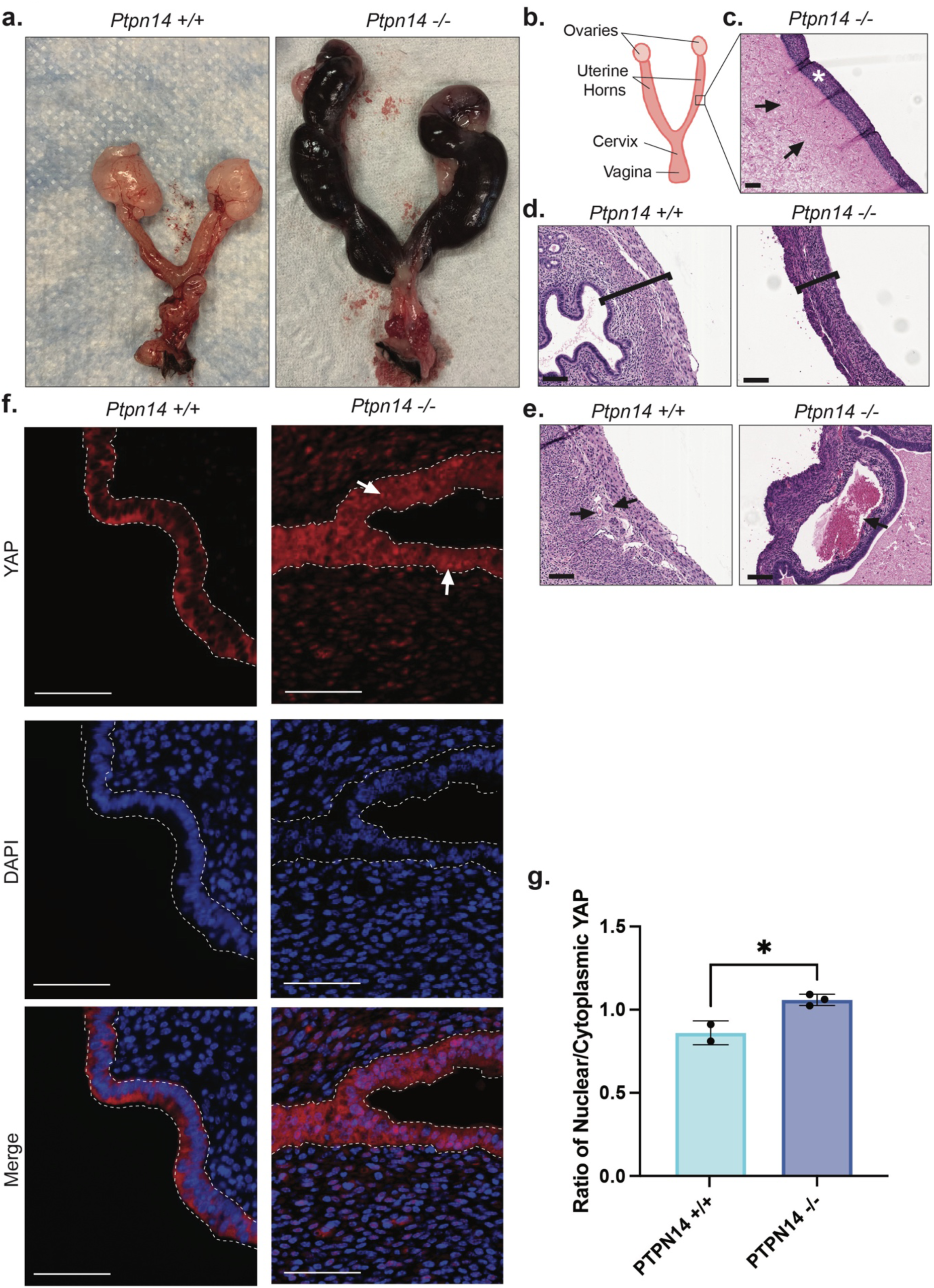
**Characterization of Hydrometra in *Ptpn14^-/-^* Mouse Uteri**. (A) Whole mount images of representative *Ptpn14^+/+^* (left) and *Ptpn14^-/-^* (right) mouse uteri. A total of 12 female mice were dissected and evaluated for hydrometra. These mice comprised those in the aging cohort (n = 8, excluding the mice that were found dead) and other mice in the colony (n = 4). These ranged in age from 171 to 499 days. (B) Diagram of the female mouse reproductive system. The black box indicates an example of the regions shown in C-F. (C) H&E image of the uterine wall and the fluid filling the uterus of a *Ptpn14^-/-^* mouse with hydrometra. The Uterine musculature is denoted by an * and the fluid is indicated with arrows. Scale Bar: 100 microns. (D) H&E images of longitudinal sections of the uterine walls of *Ptpn14^+/+^* (left) and *Ptpn14^-/-^* (right) mice. Brackets indicate the bounds of the uterine wall region, with *Ptpn14^-/-^* mice exhibiting a thinner wall due to the dramatic distension of the uterus. Scale Bar: 100 microns. (E) H&E images of normal blood vessels in wild-type mice (left) and the distended blood vessels found in *Ptpn14^-/-^* mice (right). Arrows indicate blood vessels. Scale Bar: 100 microns. (F) Representative YAP immunofluorescence staining in wild-type (n=2) and *Ptpn14*^-/-^ (n=3) mouse uteri. The white dotted lines indicate the epithelial regions counted in (F) for the quantification of the YAP staining. Scale Bar: 50 microns. White arrows indicate examples of nuclear YAP staining. (G) Quantification of the ratio of YAP nuclear to cytoplasmic localization in mouse uteri.

Because PTPN14 is known to regulate YAP/TAZ nuclear localization, we investigated YAP/TAZ localization in the epithelium of aged female mouse uteri^23^. PTPN14 deficiency led to an increase in the nuclear:cytoplasmic ratio of YAP in the epithelium of *Ptpn14*^-/-^ mouse uteri compared to wild-type mice (Figure 2F, G), indicative of enhanced YAP/TAZ signaling. During the estrous cycle, YAP/TAZ oscillate between nuclear and cytoplasmic localization^30^. While these mice were aged a year or more and should be in estropause, the increased YAP signaling observed in the knockout mice resembles that of younger mice in their estrus phase^30^. In support of the observation that PTPN14 contributes to uterine function, *Ptpn14* mRNA is highly expressed in the mouse uterus (Supplemental Figure 2).

### Ptpn14 Deficiency Negatively Impacts Vital Organs

To further understand the role of PTPN14 *in vivo*, we next analyzed *Ptpn14^-/-^* mice for abnormalities in vital organs. We thoroughly surveyed the effects of *Ptpn14* loss on these organs by assessing H&E staining of tissues from wild-type and *Ptpn14^-/-^* mice in our colony (n >= 3 per genotype for each organ), combined with observations from our aging cohort. Further investigation of our aging mouse cohort revealed that one 171-day-old female *Ptpn14^-/-^* mouse developed an abnormal-looking kidney with dramatic glomerular cysts, suggesting that loss of PTPN14 may disrupt kidney function over time (Figure 3A, B). However, we observed no further kidney abnormalities by gross inspection or histological analysis in the remaining mice in the aging cohort or in the other *Ptpn14^-/-^* mice in the colony (Figure 3C). This led us to hypothesize that the knockout mouse with kidney cysts was exposed to a unique stress or injury that revealed the lack of *Ptpn14* expression in the kidney. Because Hippo pathway inactivation can cause increases in organ size, we also assessed the weight of the kidneys of these mice relative to their total body weight (Figure 3D). These data indicate that *Ptpn14* deficiency does not affect kidney size, which is consistent with the relatively low expression of *Ptpn14* in mouse kidneys at homeostasis (Supplemental Figure 2, Figure 6A).

**Figure 3.**
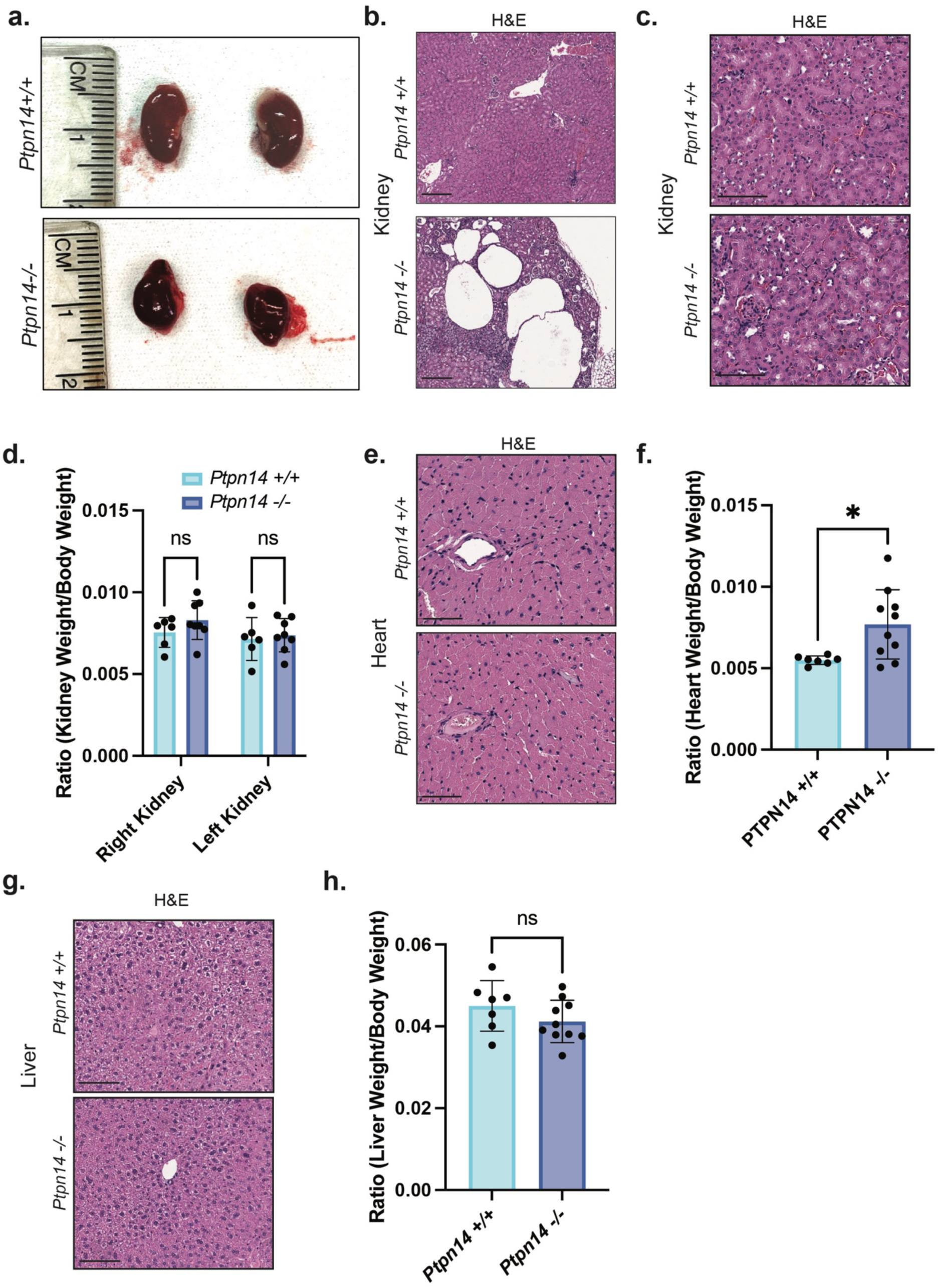
*Ptpn14* Deficiency in Organs Typically Affected in Hippo pathway Knockout Mice. (A) Whole-mount images of kidneys from a wild-type female mouse from the colony (top) and a *Ptpn14^-/-^* female mouse from the aging study (bottom). (B) H&E images of the kidneys from a *Ptpn14^+/+^* (top) mouse and the *Ptpn14^-/-^* (bottom) mouse pictured in (A), showing dramatic glomerular cysts. Scale Bar: 250 microns. (C) Representative H&E images of the kidneys from age-matched *Ptpn14^+/+^* (top) and *Ptpn14^-/-^* (bottom) mice (both male aged 100 and 98 days, respectively). Scale Bar: 100 microns. (D) The ratio of the right and left kidney weight to the body weight of *Ptpn14^+/+^* (n = 7, 5 males and 2 females) and *Ptpn14^-/-^* (n = 9, 6 males and 3 females) mice. (E) Representative H&E images of the hearts from *Ptpn14^+/+^* (top) and *Ptpn14^-/-^* (bottom) mice. Scale Bar: 50 microns. (F) The ratio of the heart weight to the body weight of *Ptpn14^+/+^* (n = 5, 5 males and 2 females) and *Ptpn14^-/-^* (n = 10, 5 males and 5 females) mice. Representative H&E images of livers from *Ptpn14^+/+^* (top) and *Ptpn14^-/-^* (bottom) mice. Scale Bar: 100 microns. (B) The ratio of the liver weight to the body weight of *Ptpn14^+/+^* (n = 7, 5 males and 2 females) and *Ptpn14^-/-^* (n = 10, 5 males and 5 females) mice. For organ weight figures, all organs measured were from mice aged over 220 days.

As previous studies reported abnormal heart growth and cardiac hyperplasia with either Hippo pathway knockout or YAP/TAZ overexpression, we next analyzed the hearts in *Ptpn14^-/-^* and wild-type control cohorts^31,32^. We measured heart weight relative to the body weight of each mouse and generated H&E sections of the hearts for histological evaluation. While histological analysis failed to show a difference in heart tissue morphology (Figure 3E), these analyses revealed that *Ptpn14^-/-^* mice had significantly larger hearts than the wild-type controls (Figure 3F). *Ptpn14* is highly expressed in multiple cell types of the mouse heart^33^ (Supplementary Figure 2,).

As hepatomegaly and hepatocellular carcinoma manifest in many Hippo pathway knockout mice, we next investigated the effects of *Ptpn14* loss in the liver^10–14,34^. Upon dissection, we observed no obvious difference between *Ptpn14^-/-^* livers and their wild-type counterparts by visual inspection and no evidence of hepatocellular carcinoma or hyperplasia in the *Ptpn14^-/-^* livers by histopathological evaluation (Figure 3G). There was also minimal difference in the liver to body weight ratios of these mice (Figure 3H). Consistent with these observations, liver cells show little to no expression of *Ptpn14* mRNA (Supplemental Figure 2). Together, these findings suggest that PTPN14 does not significantly influence liver health under basal conditions. Similarly, we detected no clear differences between *Ptpn14^-/-^* and *Ptpn14^+/+^* mice in other major organs, including the lungs, pancreata, and skin, by histological analysis (Supplemental Figure 3), despite being highly expressed in some of these tissues, such as the skin and lungs. This high-level expression suggests that PTPN14 only plays a role in these tissues under specific conditions, which may explain the lack of a clear knockout phenotype in these tissues at homeostasis (Supplemental Figure 2).

### PTPN14 Regulates Corneal Progenitor Cell Proliferation and Differentiation

One of the most prominent phenotypes observed in the *Ptpn14^-/-^* mice was the development of white eye lesions upon aging. The eyes also appeared more sunken into the skull, with swelling and hair loss in the surrounding eyelid region, suggesting irritation of the orbit. Of the 17 *Ptpn14^-/-^* mice in our aging cohort, 9 (53%) of them presented with severe eye lesions (Figure 4A). Moreover, eight out of the nine mice with eye lesions were female, suggesting that sex-specific differences may influence this phenotype. The single male that we identified had a much less severe phenotype, with no protrusion from the eye and only a subtle clouding of the cornea with cells in the epithelium that appeared slightly more proliferative. To fully characterize these eye lesions, we used knockout mice from our aging study and the rest of the colony that developed the phenotype as well as wild-type control mice from the colony. Fluorescein staining of these eyes showed extensive damage to the epithelial layer of the cornea (Figure 4B). Analysis of H&E-stained eyes revealed an irregular expansion of the basal epithelial cells in the cornea (Figure 4C, D). We confirmed this observation by co-staining for Keratin 12 (K12), a marker of corneal epithelial cells, and Ki67, a proliferation marker, which indicated that the basal epithelial cells of the cornea were indeed more proliferative in *Ptpn14^-/-^* mice than in wild-type mice (Figure 4E, F). *Ptpn14* mRNA is detected in the cornea and other cell types of the eye, supporting its role as a regulator of this tissue (Supplemental Figure 2).

**Figure 4.**
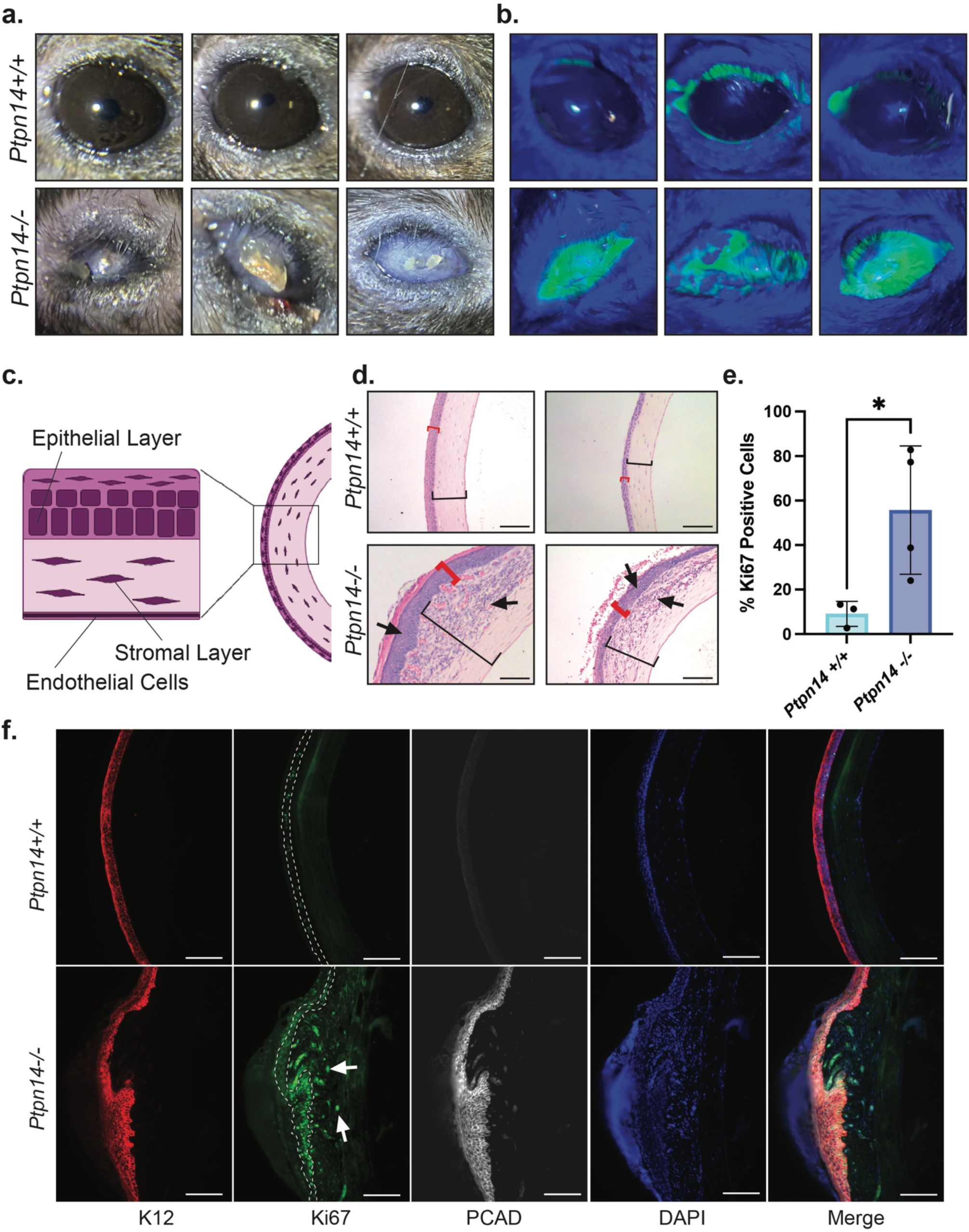
*Ptpn14*^-/-^ Mice Present with Overproliferative Corneal Cells. (A) Whole-mount images of eyes from *Ptpn14*^+/+^ (top) and *Ptpn14*^-/-^ (bottom) mice. *Ptpn14*^-/-^ mice presented with large white protrusions that extended from the center of the eye. These particular mice were not part of the aging study, although those in the aging study presented with similar lesions. (B) Fluorescein staining of eyes pictured in (A). Dramatically increased staining in *Ptpn14*^-/-^ mice indicates severe damage to the corneal epithelial layer of these eyes. (C) Diagram of the various tissue layers of the mouse cornea. (D) H&E images of *Ptpn14*^+/+^ (top) and *Ptpn14*^-/-^ (bottom) mouse corneas. Black brackets indicate the stromal layer while red brackets denote the epithelial layer. Arrows indicate regions of overproliferation in the *Ptpn14*^-/-^ mouse corneas. Mice presenting with these lesions were aged between 171 and 499 days, with a median of 219 days when the lesion formed. Scale Bar: 100 microns. (E) Quantification of Ki67 positive cells in the basal epithelial layer of the wild-type (n=3, from the colony) and *Ptpn14*^-/-^ (n=4, 2 from the aging study and 2 from the colony) mouse corneas. (F) Representative K12, Ki67, and PCAD immunofluorescence staining in wild-type (n=3) and *Ptpn14*^-/-^ (n=4) mouse corneas quantified in (E). The white dotted lines indicate the basal epithelial regions counted in (E) for the quantification of the Ki67 staining. Scale Bar: 100 microns.

Because of the stochastic nature of these eye lesions, we hypothesized that a random stress, such as a scratch, triggered hyperproliferation and an aberrant injury response that resulted in the observed lesions. Indeed, in both *Ptpn14^+/+^* and *Ptpn14^-/-^* male and female neonatal mice, the cornea developed normally, suggesting that the eye phenotypes we observed in adults were not due to developmental defects, but rather an uncontrolled injury response (Supplemental Figure 4A). In the adult mouse, the cornea is a highly regenerative organ, and its cells are in a state of constant renewal^35^. At homeostasis, the limbal epithelium, a stem cell compartment of the cornea, releases cells that slowly differentiate into mature corneal epithelial cells, which migrate toward the outer edge of the corneal epithelium and are eventually exfoliated off of the ocular surface at the end of their life (Supplementary Figure 4B)^35^. Throughout the differentiation process, these cells lose expression of Keratin 19 (K19) and P-cadherin (PCAD), severely reduce their expression of Keratin 14 (K14) and start to express K12 when they complete the transition to mature epithelial cells. This process quickens during injury, when the corneal epithelial stem cells increase their proliferation 40-fold and increase their migration to heal the corneal epithelium (Supplementary Figure 4B)^36^. During this injury response, YAP/TAZ also displays increased nuclear localization to enhance proliferation and migration^37^. As PTPN14 is a negative regulator of YAP/TAZ that also controls progenitor cell responses to injury in Drosophila, we hypothesized that PTPN14 loss and increased YAP/TAZ signaling may have disrupted the injury response in the corneal epithelium.

To assess the injury response in the *Ptpn14^-/-^* mouse corneas, we stained for markers of several known cell types in the cornea. We found that the corneal epithelial cells in the *Ptpn14^-/-^* mice expressed PCAD (Figure 4F, 5A) and K19, markers of corneal progenitor cells normally lost in mature corneal epithelial cells (Figure 5A). Notably, the cells retained high-level expression of K14, a protein highly expressed in corneal progenitor cells but that decreases its expression in mature corneal epithelial cells (Figure 5A). We also noted that the intensity of the PCAD and K19 staining correlated positively with the severity of the eye lesions, with lower expression in the apparently unaffected *Ptpn14^-/-^* eyes (Figure 5A). Furthermore, the corneal epithelium in *Ptpn14^-/-^* mice displayed increased YAP protein levels in both the cytoplasm and the nucleus compared with the corneal epithelium in wild-type mice (Figure 5B). Taken together, these data indicate a disruption in the normal differentiation process of the corneal epithelial progenitor cells and implicate YAP signaling in their increased proliferative capacity.

**Figure 5.**
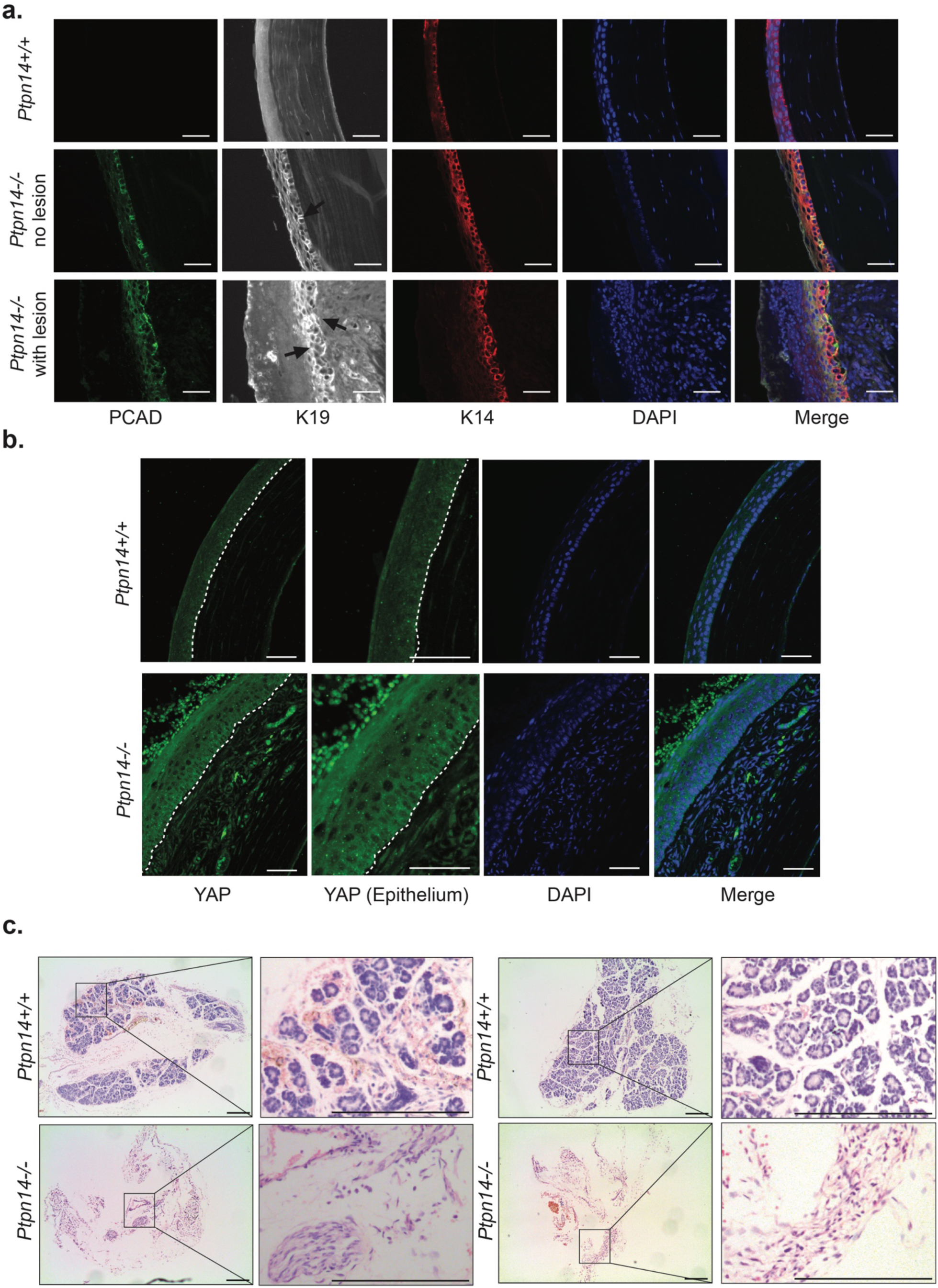
*Ptpn14* Loss Alters Corneal Epithelial Cell Maturation and Inhibits Lacrimal Gland Development. (A) PCAD, K19, and K14 immunofluorescence staining in wild-type (n=3, top row) and *Ptpn14*^-/-^ (n=3, bottom two rows) mouse corneas. (Middle) Representative image of a *Ptpn14*^-/-^ mouse corneas with no obvious eye lesion (middle row) and (Bottom) Representative image of a severe lesion. Black arrows indicate examples of regions with specific K19 staining. Scale bar: 40 microns. (B) Representative YAP immunofluorescence staining in wild-type (n=3, top) and *Ptpn14*^-/-^ (n=3, bottom) mouse corneas. Dotted line demarcates the barrier between the epithelial and stromal layers of the cornea. Scale bar: 40 microns. (C) H&E images of the lacrimal glands in neonate *Ptpn14*^+/+^ (top) and *Ptpn14*^-/-^ (bottom) mice (n= 3 for both genotypes). Scale Bar: 200 microns.

Interestingly, we also noted that the lacrimal gland, which is responsible for producing tears, was markedly smaller or absent entirely in neonate *Ptpn14^-/-^* mice compared to wild-type controls, indicating that PTPN14 is essential for normal lacrimal gland development (Figure 5C). This is consistent with the pattern of *Ptpn14* mRNA expression, which is detected in the lacrimal glands of mice (Supplemental Figure 2). While this defect likely leads to severe dry eye in the *Ptpn14^-/-^* mice, it is unlikely to be the primary stress inducing eye lesions, as the *Ptpn14^-/-^* female mice typically developed the eye lesions in only one eye, but both lacrimal glands developed aberrantly. Moreover, the severe cornea phenotype is exclusively seen in females, but analysis of our neonate cohort indicated that both male and female lacrimal glands developed improperly. Instead, we hypothesize that PTPN14 loss reprograms the corneal epithelial cells to a progenitor-like state and that injury to the corneal epithelium triggers an overproliferation of these primed progenitor cells (Supplementary Figure 4B).

### Human *PTPN14* polymorphisms are associated with a similar spectrum of phenotypes as in mice

We next sought to understand whether the many roles PTPN14 plays in maintaining the health of various mouse tissues are conserved in humans. Analysis of *PTPN14* expression in the GTEX gene expression dataset across human tissues confirmed robust expression of the RNA in the uterus, with weaker, but detectable, expression in the kidney and heart (Figure 6A), the tissues where we observed phenotypes in the *Ptpn14* knockout mice. While the eye was not included in this dataset, we found that *PTPN14* mRNA is expressed in the human lacrimal gland and cornea according to the Human Eye Transcriptome Atlas^38^. We then used data from the 53 studies in GWAS catalog to examine potential phenotypes associated with *PTPN14* variation in humans. Of the 59 associations with *PTPN14* variation identified in these GWAS studies, 9 of them were kidney-related (either glomerular filtration rate or serum creatine levels), reinforcing our hypothesis that PTPN14 plays an important role in kidney health (Figure 6B). GWAS studies also associated *PTPN14* variation with low birth weight and endometriosis (Figure 6B). Our *Ptpn14* knockout mice similarly weigh less at birth than their wild-type counterparts, and the endometriosis observed in humans could relate to the uterine abnormalities that developed in female *Ptpn14^-/-^*mice. Analysis of data from the Common Metabolic Diseases Knowledge Portal reinforced these conclusions. We noted that *PTPN14* variation was significantly associated with adult height and weight as well as birth weight, which we observed in the *Ptpn14* knockout mice (Figure 6C). The subjects in this dataset also had a significant association between *PTPN14* variation and several uterus phenotypes as well as many kidney health biomarkers, paralleling our observations in the knockout mouse uterus and kidney (Figure 6C). Furthermore*, PTPN14* variation was anecdotally associated with primary angle closure glaucoma which, when combined with our data, supports our hypothesis that PTPN14 also plays a role in eye health in humans (Figure 6C). Finally, analysis of the ExPheWas data portal associated *PTPN14* variation with traits relating to reproduction, renal health, and eye health, further reinforcing our observations in the knockout mouse (Figure 6D). Together, these data from many large human cohorts suggest that PTPN14 plays an evolutionarily-conserved role between mice and humans in many tissues and that the phenotypes in the *Ptpn14* knockout mice will provide insight into the basis for these human conditions.

**Figure 6.**
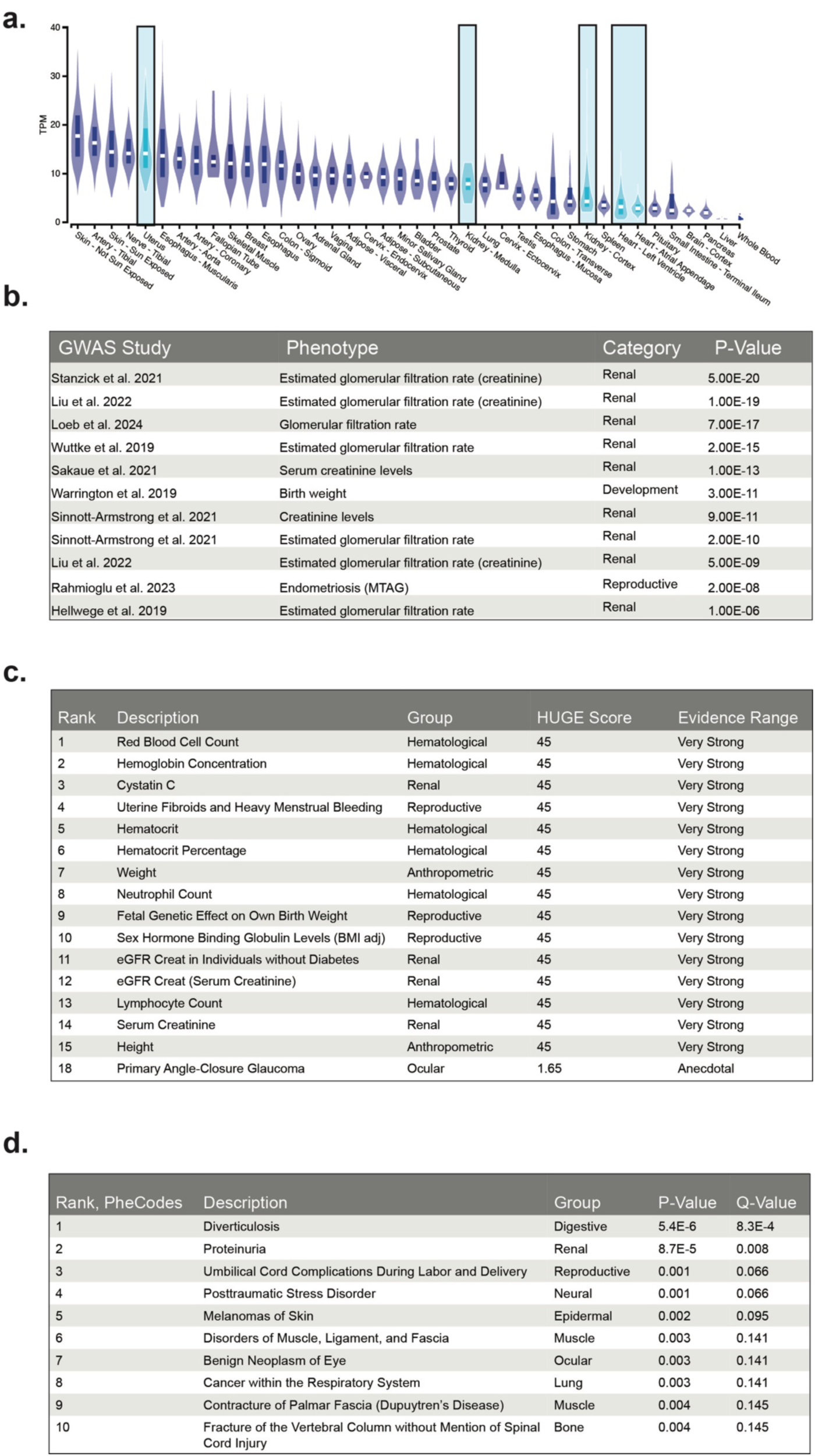
*PTPN14* Exhibits Similar Expression and Phenotypes in Humans and Mice. (A) Expression levels of human *PTPN14* across organs from the GTEX dataset. Tissues where *Ptpn14* knockout mouse phenotypes were identified are highlighted in light blue. (B) Table of GWAS studies from GWAS Catalog that identified phenotypes correlated with *PTPN14* variation that parallel phenotypes in the *Ptpn14* knockout mouse. These *PTPN14* GWAS associations were taken from a list of 59 total *PTPN14* associations identified across 49 different GWAS studies. (C) The top 15 human phenotypes associated with *PTPN14* variation from the Common Metabolic Diseases Knowledge Portal datasets. Also included is the underpowered anecdotal observations that *PTPN14* variation correlates with primary angle-closure glaucoma, as this may relate to the eye phenotype in the *Ptpn14* knockout mouse. HUGE (Human Genetic Evidence) Scores were used to evaluate the amount of genetic evidence for each association, with values >= 30 corresponding to “very strong” evidence, >= 10 “strong” evidence, >= 3 “moderate” evidence, >= “anecdotal” evidence and <= 1 representing no evidence for the given association^65^. (D) The top ten PheCode traits associated with *PTPN14* variation in humans from the ExPheWas Data portal.

## DISCUSSION

Here, we investigated the physiological roles of PTPN14 by generating *Ptpn14* knockout mice. While numerous *Ptpn14^-/-^* mice survive to adulthood, we observed significant selection against *Ptpn14^-/-^*pups prior to weaning, indicating that PTPN14 is essential for organismal viability. In contrast to other Hippo pathway protein knockout mice, which commonly fail to survive postnatally, the survival of *Ptpn14^-/-^* mice provides a unique opportunity to study the Hippo pathway across tissues in adult mice. These analyses revealed that *Ptpn14* loss is associated with a host of phenotypes in varied tissues, including the eye, uterus, heart, and kidney. Notably, by studying both male and female mice, we discovered that *Ptpn14* deficiency triggered female-specific phenotypes in the eye and uterus. Together, our analyses significantly expand our knowledge of PTPN14 and the role it plays in maintaining organ function at homeostasis.

One of the most dramatic phenotypes we observed in the *Ptpn14* knockout mice was the development of large eye lesions caused by an overproliferation of corneal epithelial cells. These lesions typically arose in only one eye, suggesting that a stochastic event, such as a scratch, elicited a subsequent injury response that was not effectively controlled. YAP/TAZ has an established role in the injury response, during which the Hippo pathway becomes inactivated to allow YAP/TAZ to initiate the growth promoting gene expression necessary for wound healing^39^. Therefore, it follows that deletion of a known negative regulator of YAP/TAZ, like PTPN14, could cause an aberrant response to injury. Indeed, YAP/TAZ has specifically been studied in corneal wound healing, where it becomes activated to promote proliferation, migration, and to disrupt cell-cell adhesion^37^. Therefore, when PTPN14 is lost, the YAP/TAZ overgrowth phenotype initiated during wound healing is sustained after the injury heals, leading to severe overproliferation of cells in the *Ptpn14* knockout mouse eyes and the consequent lesions.

While many female *Ptpn14^-/-^* mice developed eye lesions, 100% of *Ptpn14^-/-^* female mice presented with hydrometra. Although this disease is poorly understood in women, current research implicates hormonal dysregulation or obstruction or inflammation of the cervix/vagina/fallopian tubes as potential causes for hydrometra^40^. Our findings suggest that inappropriate YAP/TAZ-induced proliferation may contribute to the development of hydrometra in the *Ptpn14*^-/-^ mice. Indeed, it is known that during the estrus phase of the mouse estrous cycle, in response to hormone signaling, increased nuclear YAP/TAZ is associated with peak luminal epithelial cell proliferation^30,41,42^. YAP/TAZ also regulates injury repair in the uterus following pregnancy^41^. As proliferation during both the estrous cycle and uterine injury repair are regulated by YAP/TAZ, the increased YAP/TAZ signaling that we observe upon deletion of *Ptpn14* may reflect perturbations in the normal hormonal signaling in the uteri of *Ptpn14* knockout mice. Additionally, because obstruction of the lower reproductive track is related to hydrometra development, it is notable that PTPN14 regulates cervical epithelial function. Specifically, PTPN14 is bound and degraded by the E7 protein of high-risk Human Papilloma viruses that cause cervical cancer^43^. These observations suggest that PTPN14 restrains proliferation and serves as a tumor suppressor in the female reproductive system^43^. Indeed, HPV E7-mediated PTPN14 degradation prevents proper differentiation of primary human keratinocytes and promotes oncogenic transformation^44^. Although we failed to identify a blockage histologically, it remains possible that PTPN14 deficiency induced an aberrant proliferation of cervical epithelial cells that restricted the fluid flow out of the uterus, leading to hydrometra. Importantly, hydrometra is a predictor of cervical cancer, which, given the role of PTPN14 as tumor suppressor in the cervix, suggests the mice may have developed this cancer if we let them age longer^40^. Further investigation of the effects of PTPN14 loss during different stages of the mouse estrous cycle and at different reproductive ages will be necessary to fully establish PTPN14 as a regulator of mouse fertility, the estrous cycle, and the aging female reproductive system. To our knowledge, our studies present the first link between the Hippo pathway and hydrometra. An understanding of this process is critical, as hydrometra is estimated to affect 14.1% of postmenopausal women^45^.

Other phenotypes we observed may also relate to defective injury responses, such as the glomerular cysts that developed in the kidney. The stochastic nature of this phenotype suggests that the mouse may have experienced stress that revealed a lack of PTPN14 regulation. Indeed, a large number of GWAS studies associate human *PTPN14* variation with kidney abnormalities and diseases. Furthermore, YAP/TAZ expression increases in podocytes following injury *in vivo* and YAP/TAZ signaling plays crucial roles in the kidney’s cellular injury response^46^. With loss of PTPN14, YAP/TAZ signaling that is activated following injury may not be correctly regulated in the kidney.

While we identified many organs with phenotypes, the majority of the tissues in the *Ptpn14^-/-^* mice appeared to develop and function normally. Although PTPN14 deficiency may not affect the function of these organs, it may also be dispensable at homeostasis and play an essential a role under certain conditions, such as in response to an injury or other stress. For example, in knockout mice lacking the gene encoding AMOT, another Hippo pathway protein, in the liver, there are no apparent defects in liver function at homeostasis. However, *Amot* knockout induces YAP/TAZ-mediated overproliferation of progenitor cells following liver injury^15^. In another example, *Mst1* knockout mice display proper kidney development, but when subjected to ischaemia-reperfusion injury, exhibit reduced apoptosis in the injured cells^47^.

The sex bias in our phenotypes underscores the importance of conducting experiments in both male and female mice. For many years, female mice have been excluded from studies and, consequently, sexually dimorphic signaling pathways have been largely understudied^48^. Hydrometra was our most penetrant phenotype, affecting every female mouse. *Ptpn14^-/-^*female mice also suffered from dystocia, or difficulty giving birth, leading to compromised fertility. These phenotypes may relate to the various deleterious female reproductive phenotypes associated with *PTPN14* variation in humans (Figure 6B, C). Another predominant phenotype in female *Ptpn14^-/-^* mice was development of severe hyperproliferative eye lesions which were absent in the male mice. We originally hypothesized that these lesions might occur because PTPN14 loss disrupts lacrimal gland development and causes dry eye, but the male mice also have this defect and do not develop the severe eye lesions we observed in females. The female tendency toward this phenotype could be explained by the same factors that cause human females to experience greater frequencies of dry eye, including higher estrogen signaling and the fact that female human corneas take longer to heal from injury than male human corneas^49,50^. Because our knockout mice mirror phenotypes observed in women with *PTPN14* variation and highlight issues of great relevance to women, such as dry eye and uterine/birthing complications, our model may be useful to investigate how PTPN14 and the Hippo pathway contribute to sexual dimorphism in humans^50^.

There is very little understanding of the Hippo pathway in sex specific roles. In one recent study, the sex-specific transcription factor, VGLL3, affected Hippo pathway signaling by competing with YAP for TEAD binding^51,52^. VGLL3 initiates a transcriptional program distinct from that of YAP when bound to TEAD, which results in a female-specific autoimmune response^52,53^. The increased expression of VGLL3 in female versus male human skin could thus explain the female bias in autoimmune diseases of the skin. These findings, combined with our study, underscore the necessity of studying signaling pathways in both sexes.

Overall, the phenotypes we observe suggest that PTPN14 plays an essential, but context-specific, role in regulating YAP/TAZ signaling, with a particular importance in female biology. The many *PTPN14*-associated phenotypes detected in humans through GWAS studies, the Common Metabolic Diseases Data Portal, and ExPheWas indicate that many of the phenotypes in our knockout mice are also observed in humans. The high evolutionary conservation of PTPN14 and the consistency of *PTPN14* variation phenotypes between mice and humans suggests that our findings could provide a framework for therapeutic innovations in humans. Specifically, modulation of the Hippo pathway through either PTPN14 inhibition or blockage of YAP/TAZ signaling through TEAD inhibitors could lead to better health outcomes for women in conditions like dry eye, corneal injury, hydrometra, and other uterine complications. Future studies will further elucidate the mechanisms underlying PTPN14 function, ultimately helping to design critical therapies for improving health in women.

## ACKNOWLEDGEMENTS

We would like to thank Dr. Hong Zeng from the transgenic mouse facility for generating mice, Pauline Chu for assistance with histological samples preparation, Dr. Shengda Lin for advice on liver staining and histology, and Dr. Vivek Charu and Dr. Jonathan Scott Maltzman for examining kidney histology. We would like to thank the International Mouse Phenotyping Consortium (IMPC) for providing access to various knockout mouse model datasets on their website, www.mousephenotype.org. We also would like to thank Dr. Naeowon Kang and Dr. David Myung for generously letting us borrow equipment for imaging the mouse eyes. Finally, we would like to thank Dr. Julien Sage for careful reading of the manuscript and Dr. Stephen Montgomery for his advice on assessing human variant and GWAS data. This work was funded by the National Science Foundation Graduate Research Fellowship Program (NSF GRFP) to E.M.M. and NIH R35CA197591 to L.D.A.

## AUTHOR CONTRIBUTIONS

E.M.M. designed the conditional *Ptpn14* allele, bred and dissected the mice, designed the project, weighed the organs, generated images of the various mouse phenotypes, performed fluorescein staining of the mouse eyes, assessed pathology, wrote the manuscript, and analyzed the data. N.M. performed immunofluorescence in the eye tissues. T.T. performed immunofluorescence in the uterus tissue. H.V., B.H., W.R., and J.V. assisted with analyzing the pathology of various mouse tissue samples. M.W. assisted with dissections and weighing organs. X.Z. supervised the ocular phenotype section of this manuscript. L.D.A. provided funding, assisted with writing the manuscript, and supervised the project.

## METHODS

### Animal Studies

All mouse studies were approved and performed in compliance with the Stanford University Institutional Animal Care and Use Committee known as the Administrative Panel on Laboratory Animal Care (APLAC, protocol number 10382). Mice were housed in Stanford’s Comparative Medicine Pavilion and Shriram Center animal research facilities at 22 °C ambient temperature with 40% humidity and a 12-h light–dark cycle (7:00–19:00) in compliance with practices detailed in the National Institutes of Health and the Institutional Animal Care and Use Committee (IACUC). The Association for Assessment and Accreditation of Laboratory Animal Care provides additional accreditation to Stanford University. Mice were maintained on a pure C57BL/6 background and genotyped by PCR. Mice of both sexes were used for all experiments.

### Generation of PTPN14 Conditional and Complete Knockout Mice

To design *Ptpn14* conditional knockout mice, two LoxP sequences were inserted at Ptpn14 intron 2 and intron 3, floxing exon 3 such that deletion of exon 3 following Cre expression will result in a downstream reading frame shift with 12 nonsense amino acids and a stop codon at exon 4. These *Ptpn14^+/fl^* mice were generated using Crispr mediated genome recombination. Four guide RNAs were designed, single strand guide RNAs (sgRNAs) were purchased from Synthego (Menlo Park CA). After *in vitro* validation by DNA cutting efficiency, two sgRNAs were chosen for generating *Ptpn14^+/fl^*mouse, Ptpn14-g1 (GGCCTACGATCTC) and Ptpn14-g3 (CCAGGAGGCTTCCGGCGAAG). Single strand donor DNA containing the exon 3 locus with LoxP sites was synthesized by IDT (Iowa, USA). The donor DNA contained 150 nucleotide homologous arms on each side, exon 3 and two LoxPs inserted at the gRNA PAM site floxing exon 3. A mixture of sgRNA (10ng/µL), CAS9 protein (25ng/µL, IDT) and donor DNA (10ng/µL) were injected into C57Bl/6J pronuclear mouse zygotes. Injected embryos were then implanted into pseudo-pregnant foster mothers. A total of 215 embryos were injected, 17 pups were born, and three pups containing the LoxP sites were identified by PCR and DNA sequencing. The genotyping PCR to confirm the 5’ LoxP site used one primer upstream of the donor DNA (CATCAACGAACCCCAGCTCT), and one primer on the 3’ LoxP site (CCAGCAGCTCCATAACTTCGT). The PCR product was then sequenced to confirm the presence and sequence of the 5’LoxP site. The same PCR strategy confirmed the correct insertion of the 3’ LoxP site, with one PCR primer downstream of the donor DNA (AAGTGCAGAGTCCAACGGAG) and one primer at the 5’ LoxP site (GGCCTACGATCTCAATAACTTCGTA). This 3’ LoxP PCR product was then confirmed by sequencing. The *Ptpn14^+/fl^* founder mice were bred with C57Bl/6J for heterozygotes. The *Ptpn14^fl/fl^* mouse was achieved by breeding *Ptpn14^+/fl^* heterozygotes. Based on these analyses, we selected multiple founder mice, which we crossed to C57BL/6J mice to generate colonies of pure C57BL/6J *Ptpn14* conditional knockout mice. After the establishment of founder mice, subsequent mice were genotyped using a set of primers to identify the 5’ LoxP site (forward primer: 5’-CTTTTAGCTTGTCCCACGCTG-3’, reverse primer: 5’- AAGCAGAGGCTCGTAGCAA - 3’), the 3’ Loxp site (forward primer: 5’-CCCCCTCGAGTTCCCCATAA-3’, reverse primer: 5’-AAGTCAGGAGTCCAACGGAG-3’), the 5’ LoxP site and a region outside the donor DNA (forward primer: 5’-CGGGTGGTGTCAGATCAGAC-3’, reverse primer: 5’- CCAGCAGCTCCATAACTTCGT-3’), and the 3’ LoxP site and a region outside the donor DNA (forward primer: 5’- GGCCTACGATCTCAATAACTTCGTA-3’, reverse primer: 5’-ACATCTTTTGCCACCATGCTC-3’).

Mouse embryonic fibroblasts (MEFs) were generated from E13.5 embryos produced from a *Ptpn14^+/-^*intercross, as previously described^54^.

### Tissue Culture

MEFs were cultured in DMEM with high glucose (Gibco) supplemented with 10% fetal bovine serum and 1% penicillin-streptomycin. To test deletion of the PTPN14 protein using the floxed allele, cells were plated in 6- well plates. 24 hours after plating, cells were transduced with either Ad5-CMV-Cre (University of Iowa Viral Vector Core) or with Ad5-CMV-Empty (University of Iowa Viral Vector Core) at a multiplicity of infection of 100. 72 hours later, protein was collected for western blotting using a 2% SDS protein extraction buffer (2% SDS, 5mM Tris pH 6.8 in dH2O).

### Western Blotting

Western blotting was performed according to standard protocols. Briefly, protein quantification was performed using the Pierce BCA Protein Assay Kit (ThermoFisher). For each sample, 40 µg of protein was loaded onto a 4-12% SDS-PAGE Gel (Bio-Rad) and transferred to a PVDF membrane (Immobilon, Millipore). The membrane was blocked with a solution of 5% milk + 0.1% Tween-20 in Tris-buffered saline and then probed for either PTPN14 (1:100, Santa Cruz, sc-373766) or Alpha-tubulin (1:5000, Sigma, T6074). For secondary antibodies, blots were probed with anti-mouse or anti-rabbit HRP-conjugated secondary antibodies (1:5000, Vector Laboratories) and then developed with Clarity Western ECL Substrate (Bio-Rad) and imaged using a ChemiDoc XRS+ (Bio-Rad). Gels were analyzed using Image Lab (Bio-Rad, v.3.0).

### Immunostaining and Microscopy

Mouse tissue was dissected and collected post-mortem and fixed in formalin overnight. Fixed tissues were stored in 70% ethanol prior to embedding. Hematoxylin and Eosin staining and immunofluorescence were performed using standard protocols. For tissue immunofluorescence experiments, the following primary antibodies were used: Keratin 14 (ms-115, 1:400, Thermo Scientific), Keratin 19 (TROMA-III, 1:100, DSHB), Keratin 12 (ab18562, 1:500, Abcam), Ki67- (550609, 1:100, BD Pharmingen), Yap (sc-101199, 1:50, Santa Cruz Biotechnology), P-Cadherin (AF76, 1:1000, R&D Systems). The ratio of YAP nuclear to cytoplasmic staining was quantified using FIJI image software to measure the mean grey value of the nuclear area, which was divided by the mean grey value of the cytoplasmic area for regions of epithelial cells in the mouse uteri^55^. For evaluation of eye tissues, the eyes were carefully enucleated using a curved tweezer and are immediately fixed in 4% PFA, 20% Isopropanol, 2% Trichloroacetic acid and 2% Zinc chloride (diluted in distilled water) for one hour at room temperature, then stored in 70% ethanol at 4°C prior to processing. The tissues were washed three times with 1x PBS for 5 minutes on the rotor and eyes were placed in 15% sucrose until the tissues were settled at the bottom of the tubes and transferred to 30% sucrose overnight at 4°C. The eyes were placed inside a biopsy blocking well containing OCT at the correct orientation and were placed in -80°C until the OCT completely solidified. The eyes were sectioned on a Leica CM1850 UV-3-1 Cryostat Microtome (Leica Biosystems, US) at 10µm thickness. Antigen retrieval was performed on the sections using a heat steamer and 10mM Sodium Citrate buffer (pH 6.0) and the sections were blocked for one hour in 10% horse serum diluted in PBS. The appropriate primary antibody solution was added to the sections and was incubated overnight at 4°C. The tissues were washed three times in PBS for 5 minutes each and the appropriate secondary antibody was applied to the sections. The Slides were mounted with n-propyl gallate anti-fading reagent (P3130, Sigma-Aldrich) and imaged using a Leica DM5000- B fluorescence microscope.

