## Supplemental Material for "*Ptpn14* Knockout Mice Reveal Critical Female-Specific Roles for the Hippo Pathway"

#### Supplemental Figure 1.

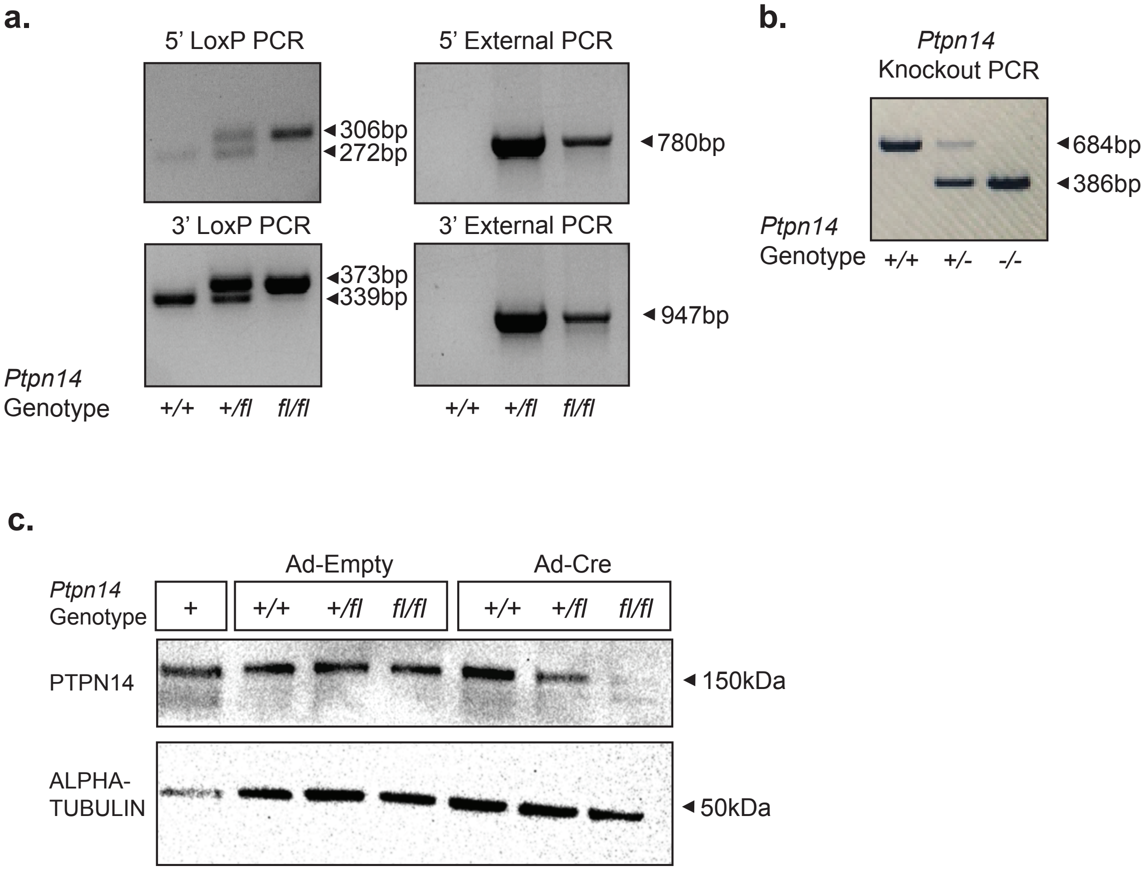


**Supplemental Figure 1. Confirming the Correct Insertion and Function of the *Ptpn14* LoxP Sites.** (A) PCR reactions performed for mouse genotyping. 5' and 3' LoxP PCRs initially confirmed the presence of both LoxP sites in each mouse. The 5' and 3' external PCRs, which amplify a product from outside the donor DNA, verified the correct location of the LoxP sites and the modified exon 3 in the genome. (B) PCR analysis of *Ptpn14*$^{+/+}$, *Ptpn14*$^{+/-}$, and *Ptpn14*$^{-/-}$ mouse DNA to confirm the successful recombination of exon 3. (C) Western blot for PTPN14 expression using MEF lines derived from a *Ptpn14*$^{+/fl}$ intercross 72 hrs after transduction with Ad-Cre or Ad-Empty. Alpha-tubulin serves as a loading control. A MEF line from a mouse where PTPN14 expression was previously detected was used as a positive control.

#### Supplemental Figure 2.
**a.**

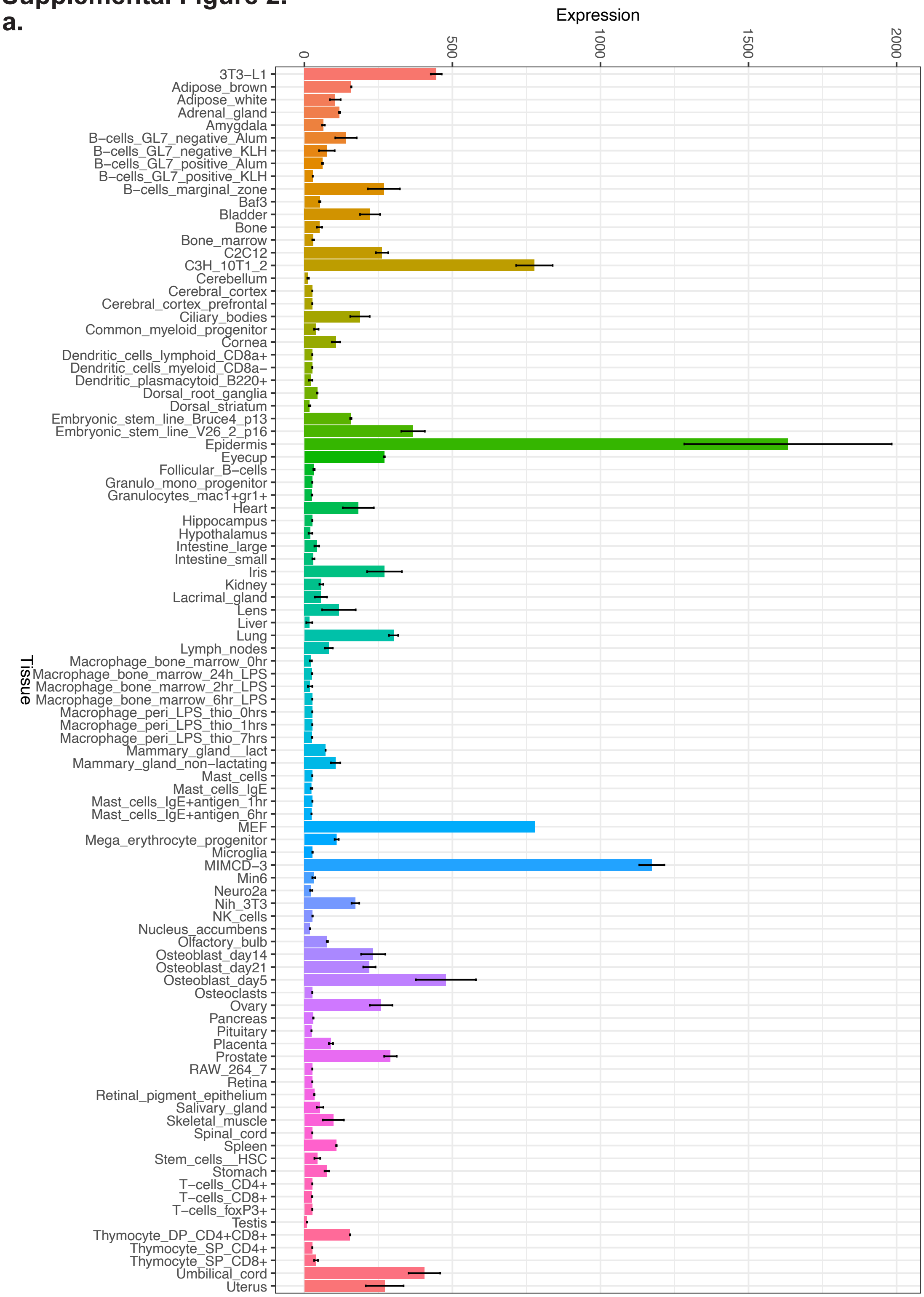


**Supplemental Figure 2.** ***Ptpn14*** **Expression Across Mouse Tissues** *Ptpn14* gene expression data across mouse tissue types from the mouse GeneAtlas dataset.

### Supplemental Figure 3.

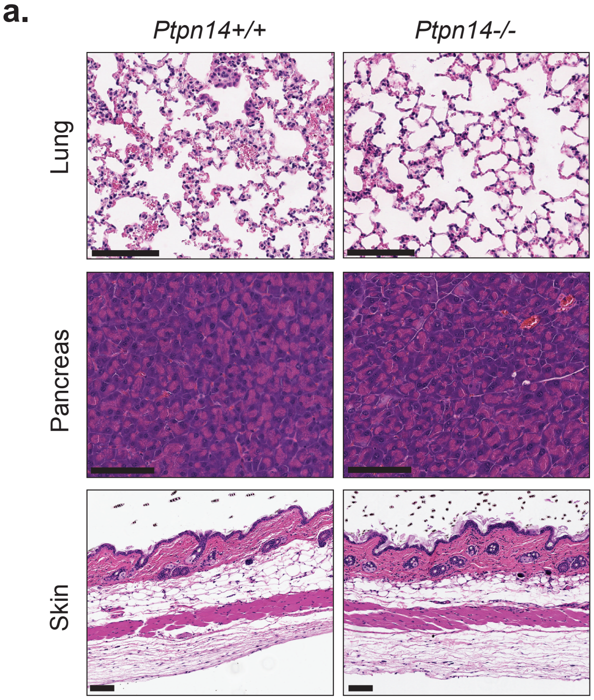


**Supplemental Figure 3. A Survey of Apparently Unaffected Organs in *Ptpn14*$^{-/-}$ mice** (A) H&E images of *Ptpn14*$^{+/+}$ (left) and *Ptpn14*$^{-/-}$ (right) mouse lung, pancreas, and skin sections. Scale Bar: 100 microns.

#### Supplemental Figure 4.

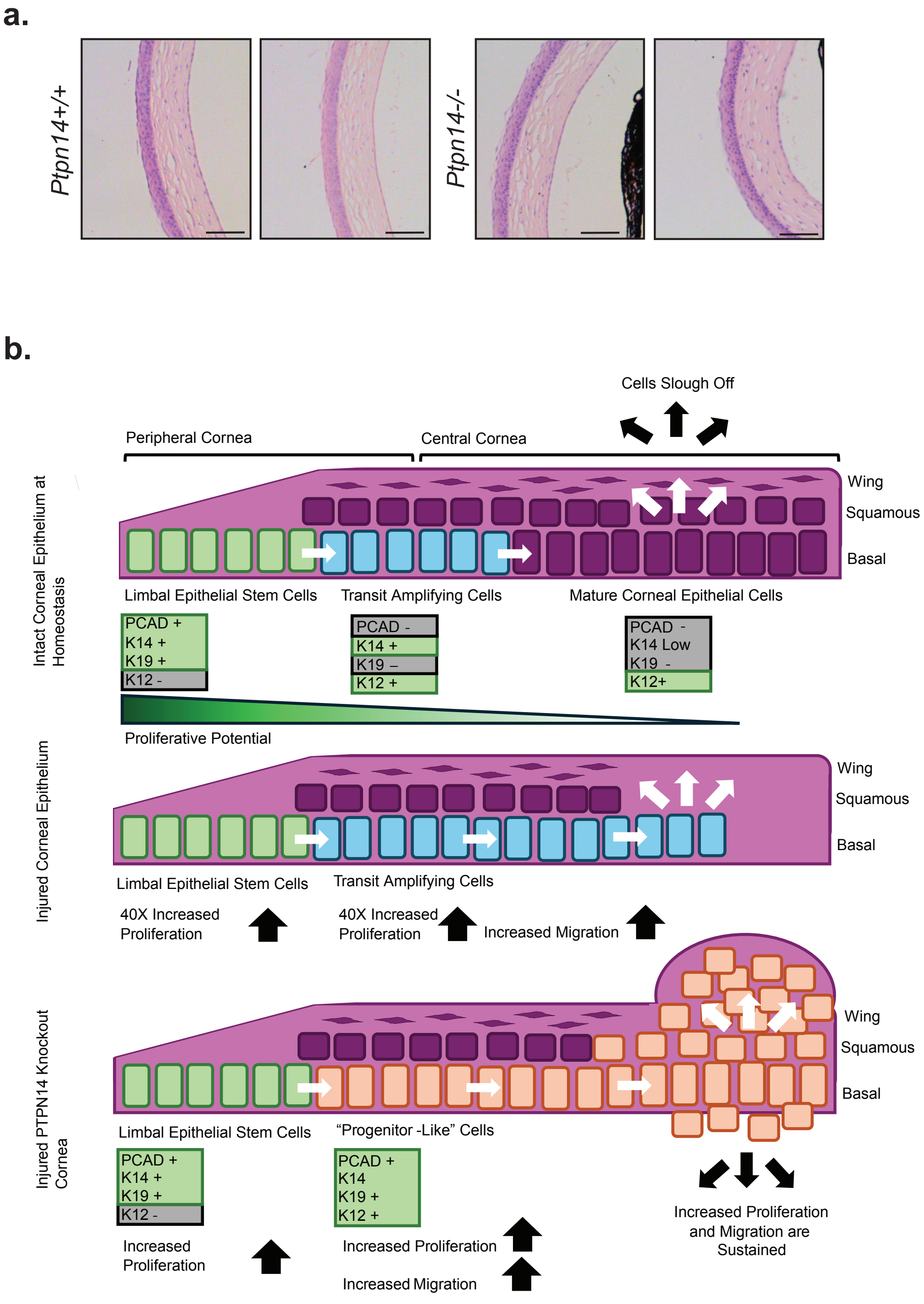


**Supplemental Figure 4. A Model for *Ptpn14* Loss in the Mouse Cornea** (A) H&E staining in neonate wild-type and *Ptpn14*$^{-/-}$ mouse corneas. Both genotypes appeared to develop normally (n=3 mice for each genotype). Scale Bar: 100 microns. (B) Diagram of mouse cornea during injury and repair. Under basal conditions, the limbal epithelial cells are the most proliferative and constantly renew the corneal epithelium. These cells reduce their proliferation and lose expression of K19 and PCAD and reduce K14 expression as they mature into corneal epithelial cells. Upon injury to the corneal epithelium, the limbal epithelial cells increase their proliferation and migration to heal the wound. In the *Ptpn14* knockout mouse, the corneal injury response is impaired and mature corneal epithelial cells maintain their increased proliferation and limbal cell gene expression.
